# Meat acquisition skill and male reproductive success in Guinea baboons

**DOI:** 10.1101/2024.09.17.613504

**Authors:** William O’Hearn, Federica Dal Pesco, Roger Mundry, Julia Fischer

**Author notes:** corresponding author Correspondence to: William J. O’Hearn, Cognitive Ethology Laboratory, German Primate Center, Kellnerweg 4, 37077 Göttingen, Germany.

## Abstract

In human foraging societies, hunting skill is often interpreted as a signal of male quality linked to his reproductive success through his ability to provision his family and community. Similarly, in some bird and insect species, males offer their mates food to indicate their quality as a provider. Among non-human primates, however, the relationship between meat sharing and reproductive success is underexplored, leaving the evolutionary origins of this phenomenon unresolved. Guinea baboons (*Papio papio*) are a promising model to investigate whether meat sharing signals male quality, since females choose their mates, have been shown to rely on males to catch and share prey, and are responsive to male foraging skill. We combined records of 109 meat-eating events with nine years of behavioural data to test whether males who more frequently acquired and shared meat with females had more females in their social units for longer. Contrary to our predictions, we found no evidence that females preferred males who acquired or shared meat more frequently, suggesting that meat acquisition does not function as a signal of male quality in Guinea baboons. One explanation may be the relatively low frequency of meat-eating events. Another is that females are less dependent on males for meat than previously reported. Our results revealed that nearly half (41%) of female meat intake originated from sources other than her unit male, including the first documented cases of female prey capture (11% of events). Thus, females likely apply other, more pertinent, criteria in mate choice.

## Introduction

In human hunter-gatherer societies, successfully catching prey can signal a male’s quality as a provider, and sharing the spoils of the hunt can broadcast that signal throughout his community (Gurven and Von Rueden 2006; Jaeggi and Gurven 2013; 2018). In such societies, meat is a highly prized food source that arrives in quantities larger than a single individual or household can consume. Consequently, the cost of sharing is small, but the benefit of receiving is significant, leading meat to be shared more widely than other foods (Hawkes, O’Connell, et al., 1991; Hill et al., 1993; Jaeggi & Gurven, 2013; Kaplan & Hill, 1985). Men who share more meat more often provide their immediate family with difficult-to-acquire nutrients, experience increased reputation within their community, and are sought after as both mates and extra-pair mating partners (Gurven 2004; Gurven and Von Rueden 2006; Hill et al. 1993; Wiessner 1996). Thus, frequent meat providers experience higher reproductive success in most hunter-gatherer societies (Smith 2004; Wiessner 1996).

While the extent of human meat sharing is unique, the practice of catching and sharing prey as a signal of male quality may have deeper evolutionary roots. Males of numerous insect (Vahed 1998) and some arachnid *Paratrechalea ornata*. Albo & Costa, 2010) and bird species (*Corvus corax*, Heinrich 1988; *Corvus monedula*, de Kort et al. 2006; *Corvus frugilegus*, Emery et al., 2007; *Sterna hirundo* (González-Solís et al. 2001); *Netta rufina*, Amat, 2000) provide prospective mates with “nuptial gifts” or “courtship feeding” to signal their quality as a provider and encourage mating opportunities, with males that share more or larger prey items generally having better reproductive outcomes. However, within the clade of our closest living relatives, non-human primates, in which we might expect to detect the evolutionary origins of human meat-sharing practices, there are no convincing examples of meat sharing as a “costly signal” of male quality (Jaeggi and Gurven 2013), nor is there substantial evidence of a relationship between male meat sharing and reproductive success (Jaeggi and Van Schaik 2011).

The bulk of the research on meat sharing has focused on chimpanzees (*Pan troglodytes*) as the only taxon that hunts cooperatively and shares meat actively (Boesch & Boesch, 1989; Watts, 2020). Studies exploring a possible link between meat sharing and reproductive success in chimpanzees have yielded mixed results. Following a meat-sharing event, male chimpanzee hunters have been reported to receive immediate copulation opportunities (Stanford 1998), delayed copulation opportunities (Achorn et al. 2023; Gomes and Boesch 2009), or none at all (Mitani and Watts 2001). Given these unclear outcomes, it may be valuable to broaden the scope beyond chimpanzees, as they are not the only primate in which meat eating occurs frequently. Indeed, chimpanzees, capuchins, and baboons are the three taxa in which meat-eating has been most often observed (Watts 2020). Both capuchins (Perry and Rose 1994) and baboons (Allan et al., 2022; Goffe & Fischer, 2016; Schreier et al., 2019; Strum, 1975; O’Hearn et al., in press) passively share meat with select individuals (tolerated theft; Stevens & Gilby, 2004). However, whether meat sharing serves as a signal of male quality in these species has not been investigated (Jaeggi and Gurven 2013; Watts 2020). Baboons, in particular, offer a valuable analogy for human hunter-gatherer diet and food practices given their shared ecological history, parallel evolution with early hominids, and, in some species, shared nested multi-level social organization (Codron et al. 2008; DeVore and Washburn 2017; Fischer et al. 2019; Patzelt et al. 2014).

In this study, we investigated Guinea baboons (*Papio papio*) as a system in which males’ ability to acquire and share meat may signal male quality to females and thus impact their reproductive success. Guinea baboons are a promising system for meat sharing as a male quality signal because: 1) females choose the male with whom they associate and mate (Fischer et al. 2017), 2) males catch prey and share meat with their females (Goffe and Fischer 2016), 3) females appear to be dependent on males to acquire meat (Goffe and Fischer 2016), and 4) females have been shown to monitor and adjust their behavior based on male foraging skill (O’Hearn et al. 2025).

Guinea baboons live in a multilevel society, at the base of which are stable ‘units’ comprised of a reproductive ‘primary male’ and one to several females with their young (Dal Pesco et al. 2022; Goffe et al. 2016). Units are nested within ‘parties,’ and parties are nested within ‘gangs’ (Fischer et al. 2017; Goffe et al. 2016; Patzelt et al. 2014). At the unit level, male-female tenure is variable – some associations last for weeks, while others last for years (Goffe et al. 2016). The choice of unit appears to be driven by females who often transfer across social levels to join a male’s unit; however, the criteria by which they choose males remain uncertain (Dal Pesco et al. 2022; Fischer et al. 2017). Once in a unit, females demonstrate high mate fidelity, with males siring 91.7% of offspring in their units, making unit size and female tenure strong indicators of male reproductive output (Dal Pesco et al. 2022). Like other baboons, Guinea baboons are eclectic omnivores (Ohrndorf et al. 2025; Zinner et al. 2021) that occasionally acquire meat from birds or small mammals (Zinner et al. 2021). We use the term “acquire meat” because Guinea baboons opportunistically catch and eat prey they encounter rather than actively “hunt” prey like humans or chimpanzees (Goffe and Fischer 2016; Hawkes, O’Connell, et al. 1991; Mitani and Watts 2001; Watts 2020). Their main prey are young bushbucks (*Tragelaphus scriptus*), which are too large for a single baboon to consume (young bushbuck: 10 – 14 kg, adult Guinea baboon: 10 – 26 kg). Meat is usually eaten by multiple individuals (O’Hearn et al., in press). A previous study of a small number of meat-eating events reported that only male Guinea baboons caught prey, but they shared it, preferentially with females in their units who ate 10-40% of the acquired meat (Goffe and Fischer 2016). Lastly, in a recent experiment, female Guinea baboons were shown to monitor their males’ foraging skills and to compete over access to them when the males alone could provide access to a high-quality food source (O’Hearn et al. 2025). Thus, female Guinea baboons appear to possess both the means and motivation to evaluate males’ meat acquisition and sharing and use that information as a signal of male quality to inform their choice of mate. We predicted that if females consider meat acquisition success and meat sharing frequency as features of male quality, males who excel at these behaviors should have 1) more associated females and 2) retain females in their units for longer.

## Methods

### Field site and study subjects

Field work for this study was carried out between April 2014 and June 2023 on the population of Guinea baboons at the field station “Centre de Recherche de Primatologie (CRP) Simenti” (13°01’34” N, 13°17’41” W), in the Niokolo-Koba National Park, Senegal. Across that period, the study population included 465 individually identified animals of all age and sex classes from both core focal parties, in which all party members were identified, and adjacent parties, in which only a few individuals were known. Baboons in the focal parties were fully habituated and individually identified by natural markings, body shape and size, and radio collars.

### Female-male associations data collection

We monitored female-male associations and unit composition in eight focal parties using observational and demographic data collected between April 2014 and June 2023. Each field day, researchers recorded information about demographic changes (i.e., births, deaths, and dispersal), took a census of the individuals seen and parties present, and recorded behavioral observations of adults and subadults using 20-minute focal follows (Altmann 1974). Behavioral observations were recorded on electronic forms using the Pendragon software (Pendragon Software Corporation, USA), which ran on cellular phones (Samsung Note 2 and Gigaset GX290). On average, we had six focal follows per individual per month, balanced across different times of day. During focal follows, researchers continuously recorded the activity of the focal animal (i.e., moving, feeding, resting, and socializing), as well as all occurrences of social behaviors, such as grooming and sitting in contact (Dal Pesco and Fischer 2025). We monitored daily female-male associations using observations of interactions between females and males (i.e., frequency of copulations, grooming bouts, contact-sit bouts, greetings, aggression, and duration of grooming and contact-sit bouts) (Dal Pesco et al. 2021; Dal Pesco and Fischer 2018). This was based on previous findings that female Guinea baboons’ primary social partner and nearly only mating partner is the male of their unit (Goffe et al. 2016)

### Meat-eating event data collection

Meat-eating events were recorded opportunistically with photographs or video recordings from video cameras (handheld Panasonic HC-X909 and GoPro Hero8) in combination with written descriptions in electronic forms using the software Pendragon. In the case of video recordings, observers verbally described the scene as it unfolded, including the identities of all adults or subadults that initially caught the prey or subsequently were transferred part or all of the carcass.

## Data analysis

### Male meat acquisition and unit size

To test our hypothesis that “males who acquire meat more often have more females in their units”, we used a generalized linear mixed model (GLMM) with a Poisson error distribution in which the response was the mode male unit size for each year, fit with the R package lme4 version 1.1-21 (Bates et al. 2015). The model had two predictors measuring males’ meat acquisition abilities: ‘prey caught’ and ‘meat acquisition’. Prey caught was the number of times a male caught prey each year. We included this term to test whether females prefer males who actually found and captured prey. Meat acquisition was defined as the total number of times a male possessed a carcass in a given year. This term was a more general measure of a male’s motivation and ability to obtain meat from his physical and social environment. The number of preys caught and meat acquired were divided by the number of meat-eating events in each male’s party in the given year to account for differences in relative availability of prey/meat. Thus, the predictors can be understood as the number of times males caught prey or acquired a carcass, given the number of opportunities to do so each year. The two predictors were z-transformed to ease model convergence and interpretation (Schielzeth 2010). We also included a categorical variable with four levels as a fixed effect in the model that was each male’s most common age category for the given year (‘mode male age’) to account for the known fact that prime-aged males have larger unit sizes (for age category definitions see CRP Simenti work manual; Dal Pesco and Fischer 2025). We included ‘male-ID’ and ‘party-ID’ as random intercepts to account for repeated observations of the same individuals and parties. Lastly, we included random slopes for prey caught and meat acquired within male- and party-IDs into the model (Barr et al. 2013; Forstmeier and Schielzeth 2011).

### Male meat acquisition and female tenure

To test our hypothesis that “females stay longer in the units of males who share meat with them more often,” we used a survival model (Cox proportional hazard model) fit with coxme version 2.2-20 (Therneau 2018). We chose this approach because it allowed us to include all male-female associations and to censor those that were still ongoing. Additionally, survival models test how the “time until an event” (i.e., the end of a relationship) is affected by the outcome of one or more variables (i.e., frequency of meat sharing), which allowed us to ask a slightly more causal form of our question: “Is the amount of time until a female leaves her primary male affected by the frequency with which he acquires meat or shares meat with her?”. The response in the model was the duration of each female-male association, from start to finish, measured in months, with an indication of whether the association was ongoing (i.e., right-censored). Some female-male dyads were already associated when data collection began in 2014 (i.e., left-censored); therefore, we regarded all associations as “minimum unit tenures,” as we did not know the extent of these associations before 2014. The main predictors were ‘meat possessions’, i.e., the frequency with which each primary male possessed a carcass, and ‘meat shared’, i.e., the frequency with which they shared meat with the female during their association. To account for observation effort, both frequencies were calculated by dividing counts of meat possession and sharing by the number of days the male was observed during the male-female association. Possession and sharing frequencies were z-transformed to ease model convergence (Schielzeth 2010). We also included whether or not a male was prime-aged (yes, no; for age categories see Simenti work manual: (Dal Pesco and Fischer 2025) as a fixed effect to control for prime-aged males’ tendency to have larger units. We used a binary measure of male age to facilitate model convergence: the dataset contained very few datapoints in certain age categories, which would have made it difficult for the model to estimate the overall effect of age. Similarly, we included ‘mode female age’ in the model, as older females would necessarily have had the opportunity to accumulate longer unit tenures than younger females. We also included ‘female-ID’ and ‘male-ID’ as random intercepts to account for repeated observations of the same individuals in different dyads. Random slopes were not included because they were not theoretically identifiable for any of the predictor-grouping pairs (Barr et al. 2013; Forstmeier and Schielzeth 2011).

### Model Validation

Before fitting our models, we inspected all quantitative predictors for roughly symmetric distributions. After fitting the models as an overall test of our main predictors and to avoid “cryptic multiple testing”, we conducted full-null model comparisons (Forstmeier and Schielzeth 2011), wherein the null model lacked the primary predictors in the fixed effects part but was otherwise identical to the full model. The comparison was based on a likelihood ratio test (Dobson 2002). After fitting the Poisson model, we checked the mean-variance assumption by examining the dispersion parameter, which did not deviate much from one (dispersion parameter = 0.621). We determined Variance Inflation Factors (VIFs) using the vif function of the car package (version 3.0-3). Assessment of Variance Inflation Factors did not reveal any collinearity issues (max VIF = 1.25) (Quinn and Keough 2023). We assessed model stability on the level of the estimated coefficients by excluding levels of the random effects factors one at a time using a function written by R.M. (Nieuwenhuis 2012). The models were all found to be suitably stable. For the Poisson model we determined 95% confidence intervals for model estimates and fitted values using a parametric bootstrap (N = 1,000 bootstraps) with the function bootMer of the lme4 package. We fitted the models in R (version 4.2.0; R Core Team 2022).

## Results

Between April 2014 and June 2023, we recorded 109 meat-eating events at a rate of approximately 1.12 per month or 2.79 events per 100 hours of observation. The events comprised of 320 meat transfers, in which 111 individuals from 13 parties participated as either possessors or recipients. In 57/64 of events in which the prey-catcher was identified, the prey was caught by males (Figure 1A). In the remaining seven events (11%), the prey was caught by females. This is the first reported observation of female prey capture in this species. Females were recipients in 140/320 transfers, 69 of which came from her primary male, and 14 others came from females in her unit originating with prey caught by the primary male (Figure 1B). Thus, females’ main source of meat was their primary male (83/140; 59%). Of the remaining 57 transfers (41%) that females received, 43 came from males other than their primary male, and 14 came from females whose meat originated from outside their unit (Figure 1B).

**Figure 1.**
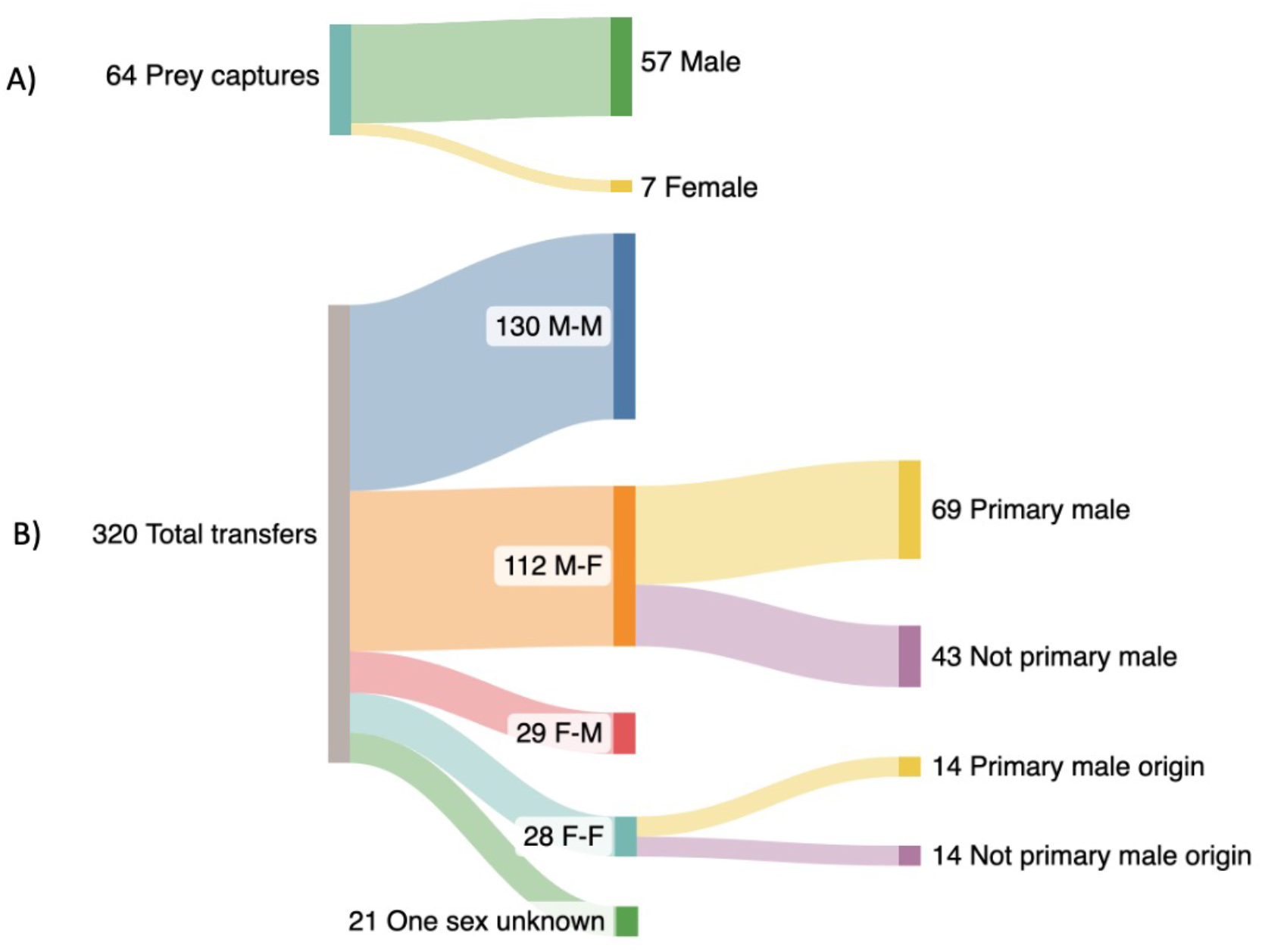
Shows two Sankey diagrams of A) prey capture events split by the sex of individual that initially caught the prey, and B) meat transfers split by the sex of possessor and recipient and in the case of female recipients, whether the meat originated from the her primary male

### Male meat acquisition and unit size

The model to assess the link between male meat acquisition and unit size included 235 observations of 57 males across eight parties with a median unit size of one (range: 0-7). The number of males that caught prey at least once was 19 (range: 1-3 times per year) and the number that acquired meat at least once was 34 (range: 1-8 times per year). The full-null model comparison was not significant (χ^2^ = 2.41, df = 2, *P* = 0.299). Thus, we found no significant relationship between the number of associated females in a male’s unit and the number of times a male caught prey (Estimate = −0.232, SE = 0.072, *P* = 0.747; Table 1) or possessed a carcass (Estimate = 0.113, SE = 0.074, *P* = 0.129; Table 1). In other words, males who caught prey (Figure 2A) or acquired meat more often (Figure 2B) did not have larger unit sizes than males who caught prey or acquired meat less often. The largest effect size of factors in the model was male age, with early and mid-prime males likely to have larger numbers of associated females in their units than sub-adult males, late-prime males, and old males (Table 1).

**Table 1.**
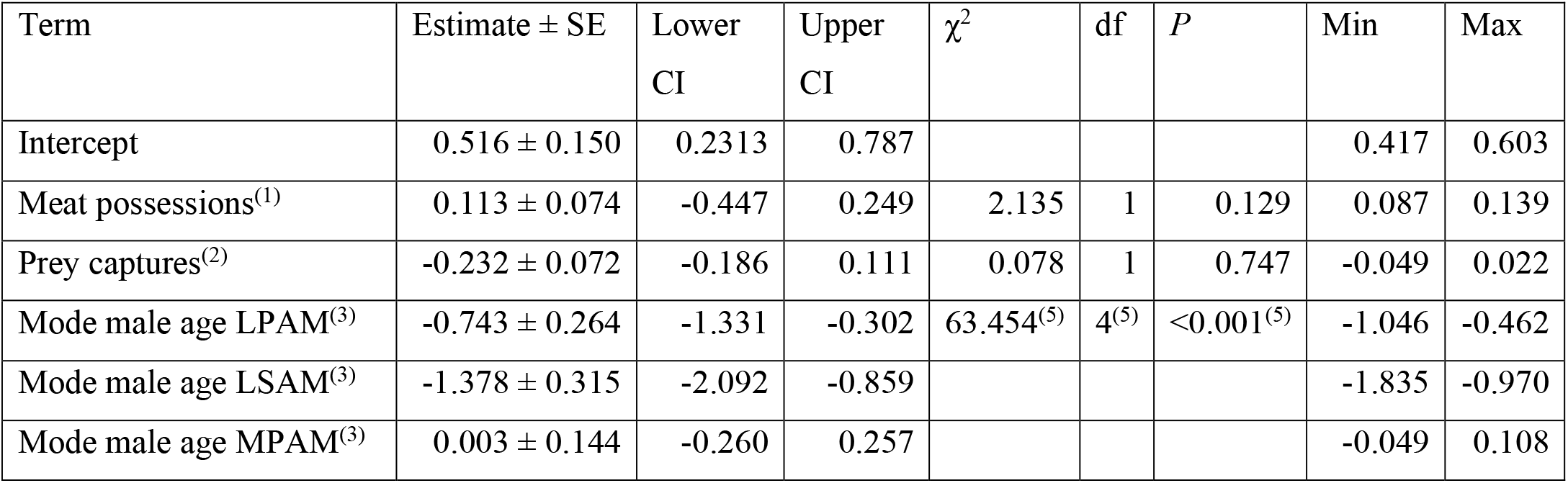
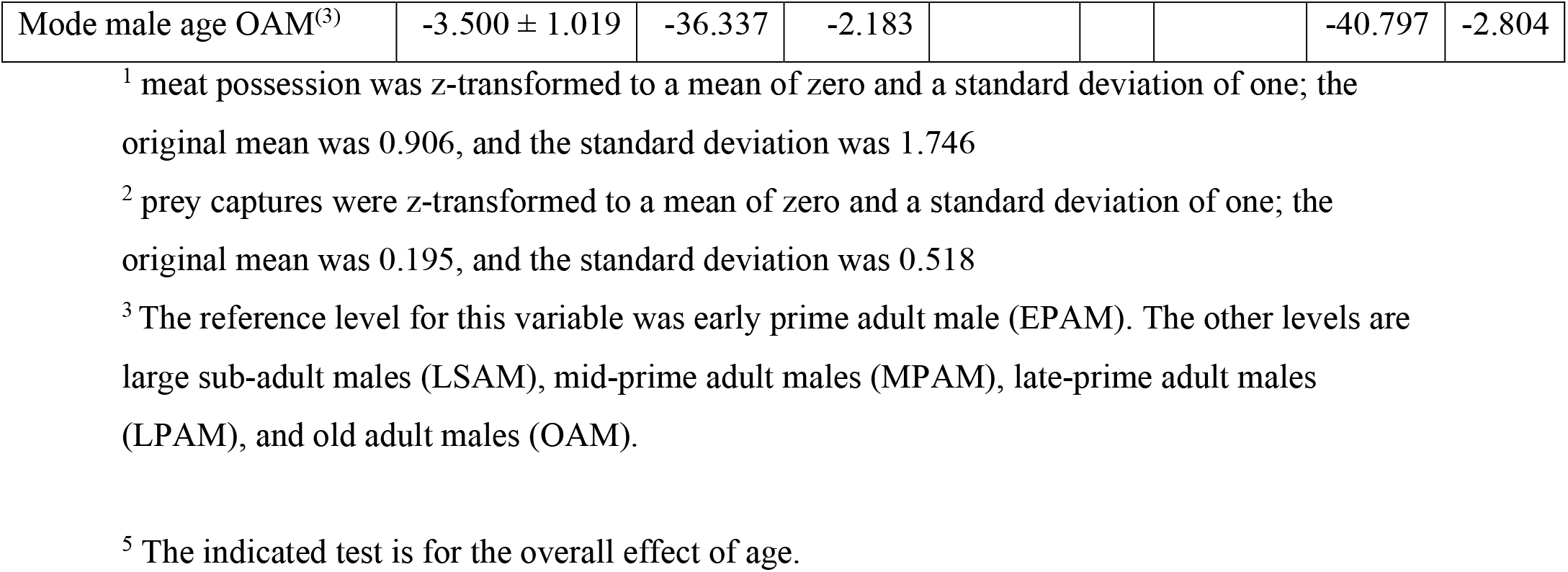
GLMM model output of male unit size predicted by meat possession and prey capture frequency.

| Term | Estimate ± SE | Lower CI | Upper CI | $\chi^2$ | df | P | Min | Max |
| --- | --- | --- | --- | --- | --- | --- | --- | --- |
| Intercept | 0.516 ± 0.150 | 0.2313 | 0.787 | | | | 0.417 | 0.603 |
| Meat possessions<sup>(1)</sup> | 0.113 ± 0.074 | -0.447 | 0.249 | 2.135 | 1 | 0.129 | 0.087 | 0.139 |
| Prey captures<sup>(2)</sup> | -0.232 ± 0.072 | -0.186 | 0.111 | 0.078 | 1 | 0.747 | -0.049 | 0.022 |
| Mode male age LPAM<sup>(3)</sup> | -0.743 ± 0.264 | -1.331 | -0.302 | 63.454<sup>(5)</sup> | 4<sup>(5)</sup> | <0.001<sup>(5)</sup> | -1.046 | -0.462 |
| Mode male age LSAM<sup>(3)</sup> | -1.378 ± 0.315 | -2.092 | -0.859 | | | | -1.835 | -0.970 |
| Mode male age MPAM<sup>(3)</sup> | 0.003 ± 0.144 | -0.260 | 0.257 | | | | -0.049 | 0.108 |
| Mode male age OAM<sup>(3)</sup> | -3.500 ± 1.019 | -36.337 | -2.183 |  |  |  | -40.797 | -2.804 |

<sup>1</sup> meat possession was z-transformed to a mean of zero and a standard deviation of one; the original mean was 0.906, and the standard deviation was 1.746
<sup>2</sup> prey captures were z-transformed to a mean of zero and a standard deviation of one; the original mean was 0.195, and the standard deviation was 0.518
<sup>3</sup> The reference level for this variable was early prime adult male (EPAM). The other levels are large sub-adult males (LSAM), mid-prime adult males (MPAM), late-prime adult males (LPAM), and old adult males (OAM).
<sup>5</sup> The indicated test is for the overall effect of age.

**Figure 2.**
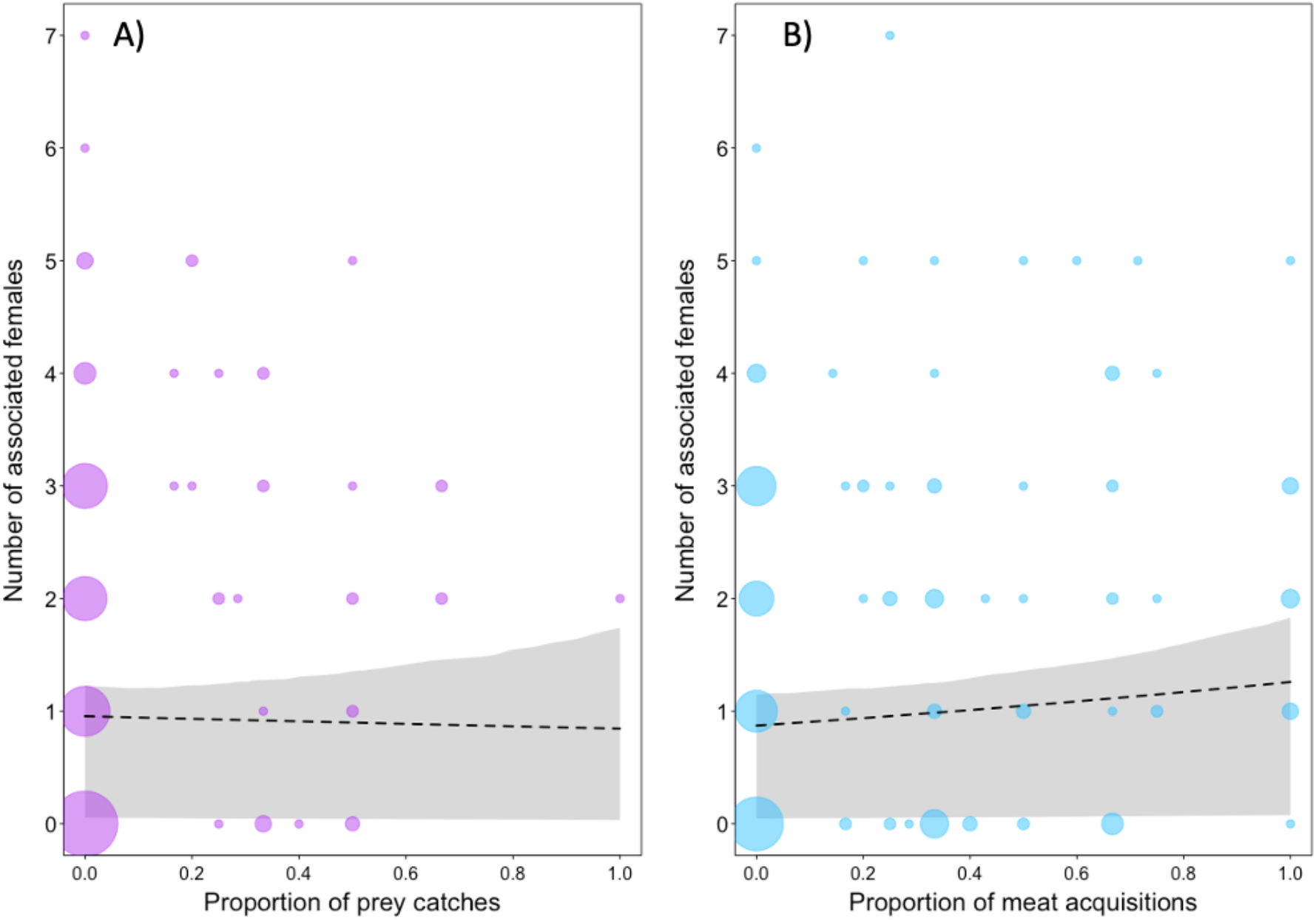
Relationship between the number of associated females (mode per year) and A) the proportion of times males captured the prey item or B) acquired meat from all the meat-eating events in their party each year. There was no evidence of a relationship between prey captures (GLMM: n = 235, p = 0.747) or meat acquisition (GLMM: n = 235, p = 0.129) and the number of associated females. Points represent the males in a given year (2014-2023), and the area of the points shows the number of males that have the same yearly value [range: A) 1-64, B) 1-43. The dashed line depicts the fitted model and the grey polygon with the bootstrapped 95% confidence intervals, with all other predictors being centered.

### Male meat acquisition and female tenure

The model examining the link between male meat acquisition and sharing and female tenure comprised 236 dyads, formed by 99 females and 54 males, with a median association duration of 269 days (range: 1-2967 days). Of those dyads, there were 55 in which the male acquired meat at least once (range: 1-12) and 28 dyads in which males shared meat at least once (range: 1-12). The full-null model comparison was not significant (χ^2^ = 3.08, df = 2, *P* = 0.21). Consequently, we found no significant relationship between the duration of a female’s tenure and the number of times her male acquired meat (Estimate = −0.103, SE = 0.186, *P* = 0.577; Table 2; Figure 3A) or shared meat with her (Estimate = −0.157, SE = 0.195, *P* = 0.419; Table 2; Figure 3B). As in the unit size model above, we found that prime-aged males were more successful in maintaining a unit, as females associated with them for longer than with non-prime-aged males (Estimate = 2.971, standard error = 1.152).

**Table 2.** Survival model output for female unit tenure predicted by male meat possession and sharing frequency.

| Term | Estimate ± SE | $\chi^2$ | df | P | Min | Max |
| --- | --- | --- | --- | --- | --- | --- |
| Male sharing events<sup>(1)</sup> | -0.154 ± 0.192 | 0.236 | 1 | 0.626 | -0.291 | 0.071 |
| Male meat possessions<sup>(2)</sup> | -0.089 ± 0.177 | 5.011 | 1 | 0.251 | -0.226 | 0.040 |
| Mode male age NP<sup>(3)</sup> | 2.975 ± 1.150 | 5.223 | 1 | 0.009 | 2.516 | 3.675 |
| Mode female age OAF<sup>(4)</sup> | 0.765 ± 1.304 | 6.645 | 3 | 0.156 | -22.851 | 1.243 |
| Mode female age SAF<sup>(4)</sup> | 1.145 ± 0.504 | | | | 0.992 | 1.352 |
| Mode female age YAF<sup>(4)</sup> | 0.562 ± 0.534 | | | | 0.341 | 0.760 |

<sup>1</sup> mean = 0.207 and SD = 0.779
<sup>2</sup> mean = 0.648 and SD = 1.68
<sup>3</sup> NP refers to non-prime aged. The reference level for this variable is prime-aged.
<sup>4</sup> The reference level for this variable is mature adult females (MAF). The other levels are sub-adult females (SAF), young adult females (YAF), and old adult females (OAF). The indicated significance test is for the overall effect of male age.

**Figure 3.**
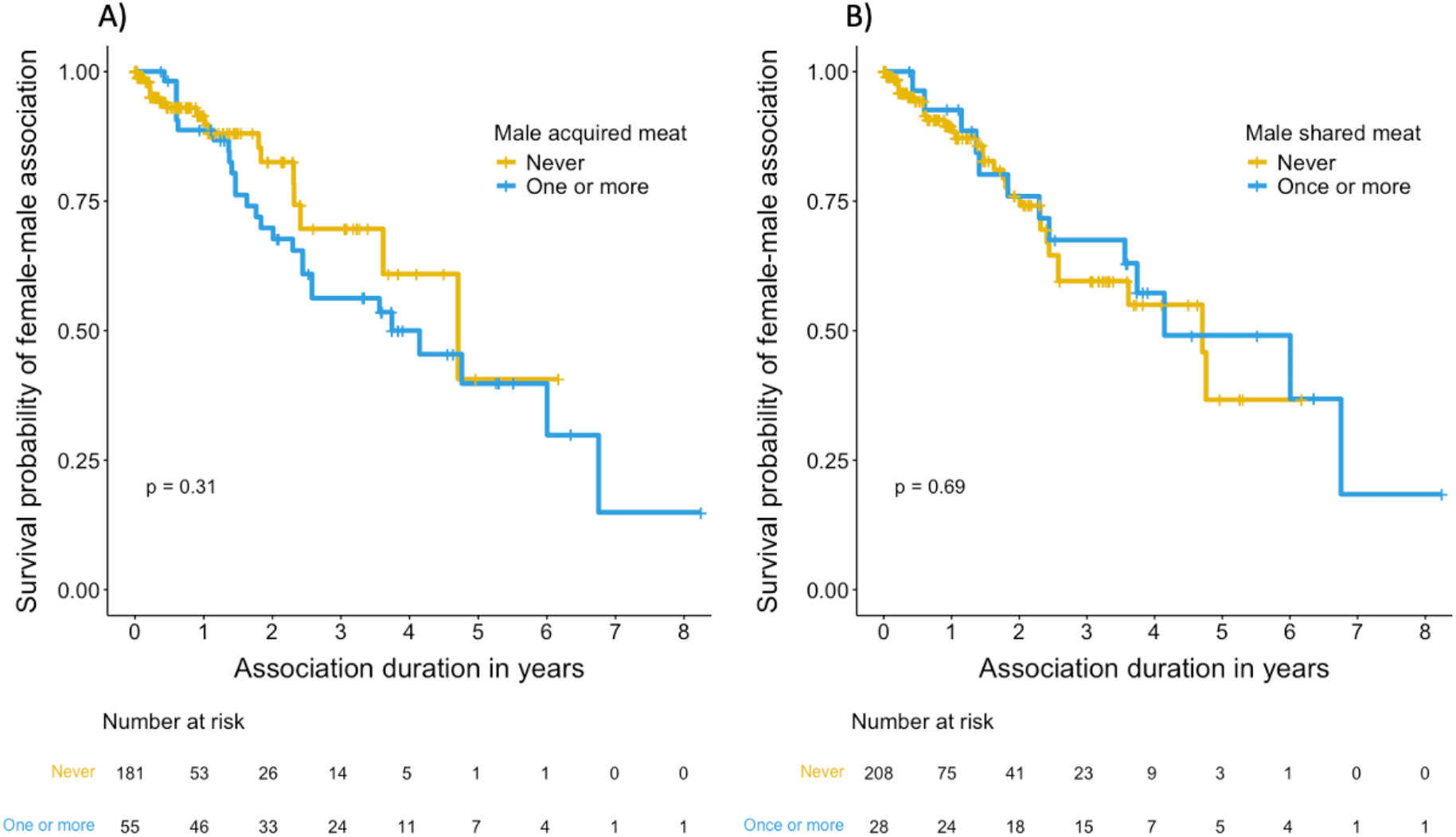
Kaplan-Meier survival curves for the effects of males A) acquiring meat and B) sharing meat with their female on the ‘survival’ of the male-female association. Separate lines are for males who never acquired or shared meat (blue) and males who acquired or shared meat at least once (yellow). This figure is a simplified to illustrate the result. The model included all possible values of meat acquisition and sharing. Risk tables beneath the plots give the number of female-male associations that ‘survived’ to each yearly increment.

## Discussion

We found no evidence that female Guinea baboons preferred to join or remain in the units of males who acquired or shared meat more often. Thus, prey capturing and meat sharing are unlikely to be signals of male quality in Guinea baboons. Although females increase affiliative interactions with their primary males to access desirable food in his possession (O’Hearn et al. 2025), they do not appear to use males’ meat acquisition or sharing skills to choose mates, suggesting that these traits may not be relevant for female reproductive decisions.

The simplest explanation for female Guinea baboons not preferring males based on their record of acquiring meat is that meat-eating events are relatively infrequent in the species. Rates of prey capture in Guinea baboons (2.79 prey per 100 hours, 1.12 prey per month) are comparable to other species of baboons (*Papio ursinus*, 1.27 prey event per 100 hours; *Papio hamadryas*: 0.1 – 2.8 prey per 100 hours; *Papio cynocephalus*: 1.78 prey per 100 hours; *Papio anubis*: 4.54 prey per 100 hours, Hausfater, 1976; Kummer, 1968; Schreier et al., 2019; Watts, 2020), and capuchins (*Cebus capucinus*: 5.35 prey per 100 hours; *Sapajus libidinosus*: 0.3 prey per 100 hours, Watts, 2020). However, prey capture rates are generally lower than capture rates of chimpanzees (Kanyawara – Ngogo: 0.8 – 7.6 per month, Watts, 2020) and far below the weekly hunts of many human hunter-gatherer societies (Hill et al. 1993; Kishigami 2000; Naylor et al. 2021). The infrequent occurrence of meat-eating events could mean that the males’ ability to acquire meat has little impact on female fitness or least less impact than other traits which females might use as criteria when choosing a primary male. Male behavioral tendencies such as their willingness to care for offspring (Rosenbaum et al. 2018), degree of tolerance toward unit females (Goffe et al. 2016), or general attentiveness to social or external threats likely outweigh the impact of meat provisioning, as they more consistently and directly impact females’ reproductive output and safety. Additionally, other features of the unit, such as the number and identities of other females and their offspring, may also influence female unit choice and tenure. Females may avoid units with antagonistic females or prefer those with existing affiliative partners, as seen in female mountain gorillas (*Gorilla beringei beringei*) who preferred to join a male’s harem when it contained females with whom they were already familiar (Martignac et al. 2025).

Another reason male meat acquisition may not impact female partner preference could be that females have more pathways to access meat than previously reported (Goffe and Fischer 2016). In our expanded data set covering nine years of meat-eating events, primary males were indeed the main source of meat for their unit females (59%). However, nearly half of the meat consumed by females came from sources other than their primary male (Figure 1). Additionally, we report the first observations of female prey capture in this species (11% of prey captures), which is consistent with evidence of prey capture by both sexes in other baboon species (Allan et al. 2022; Schreier et al. 2019; Strum 1975). Together, these findings indicate that female Guinea baboons are less dependent on males for meat access than was previously thought, which could further explain the lack of mate preference for males that acquire more meat.

In conclusion, meat acquisition and sharing do not appear to function as a signal of male quality in Guinea baboons. This finding is consistent with the wider pattern in primates that male-to-female meat sharing is not clearly associated with increased reproductive opportunities for males (Jaeggi & Gurven, 2013; Jaeggi & Van Schaik, 2011; but see Gomes & Boesch, 2009). Our results thus underscore the difference between the role of meat sharing in human foraging groups and non-human primates. Given the evidence, the human hunting and meat-sharing patterns may not stem from the evolutionary past shared with other primates. Instead, it might reflect the more recent development of the human foraging niche, which relies on difficult-to-acquire, nutrient-dense resources that promote widespread sharing (Gurven 2004; Kaplan and Gurven 2005).At the same time, our findings raise important questions specific to Guinea baboons. If meat acquisition and sharing are not central to male quality assessment, what criteria do females rely on when choosing males? Similarly, if the capacity to observe and evaluate others’ foraging skills did not evolve in this system to inform mate choice, then what selective pressure brought it about?

## Ethics declaration

This research was conducted within the regulations set by Senegalese authorities and the guidelines for the ethical treatment of nonhuman animals set down by the Association for the Study of Animal Behaviour (ASAB Ethical Committee/ABS Animal Care Committee, 2023).

### Acknowledgements

We want to thank the Direction des Parcs Nationaux (DNP) and the Ministère de l’Environnement et de la Protection de la Nature (MEPN) du Sénégal and the park conservateurs of the last 9 years for approval to conduct this study in the Parc National du Niokolo-Koba (PNNK). We are grateful to the CRP team of the last nine years for their help with the data collection. This research was funded by the Deutsche Forschungsgemeinschaft (DFG, German Research Foundation) – Project-ID 454648639 SFB 1528 „Cognition of Interaction” and Project-ID 254142454/GRK 2070 “Understanding Social Relationships”. Support by the Leibniz ScienceCampus “Primate Cognition” (Audacity Fund AF2020_03) is gratefully acknowledged.

## Author contributions

W.O., F.D.P., and J.F. developed the study’s concept and design; W.O. extracted data from the meat-eating event records; F.D.P. curated and extracted the demographic and behavioral data, and assisted with data preparation; W.O., F.D.P., and R.M. designed and conducted the analysis with supervision from J.F.; W.O. drafted the manuscript; and W.O., F.D.P., R.M., and J.F. reviewed the manuscript and provided final approval of the submitted version.

## Data and code availability

The data and code associated with the analysis of this project can be found at https://osf.io/9thk4/

